# Using a Human Circulation Mathematical Model to Simulate the Effects of Hemodialysis and Therapeutic Hypothermia

**DOI:** 10.1101/2021.02.11.430843

**Authors:** Jermiah J. Joseph, Timothy J. Hunter, Clara Sun, Daniel Goldman, Sanjay R. Kharche, Christopher W. McIntyre

## Abstract

**Background:** The human blood circulation is an intricate process regulated by multiple biophysical factors. Our patients often suffer from renal disease and atrial fibrillation, and are given treatments such as therapeutic hypothermia, exercise, and hemodialysis. In this work, a hemodynamic mathematical model of human circulation coupled to a representative dialysis machine is developed and used to explore causal mechanisms of our recent clinical observations.

**Methods:** An ordinary differential equation model consisting of human whole body circulation, baroreflex control, and a hemodialysis machine was implemented. Experimentally informed parameter alterations were used to implement hemodialysis and therapeutic hypothermia. By means of parameter perturbation, four model populations encompassing baseline, dialysed, hypothermia treated, and simultaneous dialysed with hypothermia were generated. In model populations, multiple conditions including atrial fibrillation, exercise, and renal failure were simulated. The effects of all conditions on clinically relevant non-invasive measurables such as heart rate and blood pressure were quantified. A parameter sensitivity analysis was implemented to rank model output influencing parameters in the presented model.

**Results:** Results were interpreted as alterations of the respective populations mean values and standard deviations of the clinical measurables, both in relation to the baseline population. A clinical measurable’s smaller standard deviation (in comparison to baseline population) was interpreted as a stronger association between a given clinical measure and the corresponding underlying process, which may permit the use of deducing one by observation of the other.

The modelled dialysis was observed to increase systolic blood pressure, vessel shear, and heart rate. Therapeutic hypothermia was observed to reduce blood pressure as well as the intra-population standard deviation (heterogeneity) of blood flow in the large (aorta) and small (kidney) vasculature. Therapeutic hypothermia reduced shear in vessels, suggesting a potential benefit with respect to endothelial dysfunction and maintenance of microcirculatory blood flow. The action of therapeutic hypothermia under conditions such as atrial fibrillation, exercise, and renal failure was to reduce total blood flow, which was applicable in all simulated populations. Therapeutic hypothermia did not affect the dialysis function, but exercise improved the efficacy of dialysis by facilitating water removal.

**Conclusions:** This study illuminates some mechanisms of action for therapeutic hypothermia. It also suggests clinical measurables that may be used as surrogates to diagnose underlying diseases such as atrial fibrillation.

## 1. Introduction

### 1.1 Clinical motivation

Chronic kidney disease affects a significant proportion of the Canadian and worldwide elderly patient population (Hill *et al.*, 2016). Hemodialysis is a repetitive treatment that prolongs the lives of the patients while simultaneously causing deleterious side effects (McIntyre, 2010). Hemodialysis is known to promote cardiac stunning (Burton *et al.*, 2009), causes accumulative ischemic damage to multiple organs (Marants *et al.*, 2019; Qirjazi *et al.*, 2020), and alters cardiac blood flow (Kharche *et al.*, 2018). Dialysis is also known to promote cardiac arrhythmogenicity (Burton *et al.*, 2007). Our recent clinical-imaging-modelling study strongly suggests that dialysis is a causal factor in augmenting organ level blood flow heterogeneity (Kharche *et al.*, 2018), altered hepatic and renal blood flow (Marants *et al.*, 2019), and affects cerebral circulation. In addition, dialysis is a resource intensive treatment that is a significant financial burden on our health care systems (Manns *et al.*, 2007). Treatments such as therapeutic hypothermia may reduce the severity of the dialysis induced side effects as seen in large scale clinical studies (Al-Jaishi *et al.*, 2020), as well as observed in animal models (Yeung *et al.*, 2011). An augmented understanding of the mechanisms of hemodialysis induced side effects and their treatment using therapeutic hypothermia and intra-dialectic exercise (i.e. exercise provided during a dialysis session (Penny *et al.*, 2019)) will potentially contribute towards accelerating the translation of our clinical research developments to clinical practice.

### 1.2 Overview of past modelling

Blood flow modelling is an advanced field that is now routinely used to complement clinical research. Whereas multi-scale models are increasingly being used to quantitatively support clinical decision making (see e.g. (Shang *et al.*, 2019)), this study relies upon lumped parameter (0D) modelling.

A human circulation model (Shi *et al.*, 2007; Shi *et al.*, 2011) was used in our recent study where arterial (aortic) stiffening was found to be the prime cause of hypertension (Altamirano-Diaz *et al.*, 2019). A detailed and calibrated whole body model (Heldt *et al.*, 2002; Heldt *et al.*, 2010) has been deployed widely, including to predict the consequences of orthostatic stress and exercise (Diaz-Artiles *et al.*, 2019). The relative contributions of dialyzer membrane related convection and diffusion have been assessed using steady state hemodynamic modelling (Pallone *et al.*, 1989). A spatially extended description of bicarbonate dialysis has been used to illustrate the potential personalization of models (Annan, 2012), while others have evaluated the benefit of citrate in calcium dialysate profiling (Aniort *et al.*, 2017) or dialysate sodium profiling (Coli *et al.*, 1998). The primary end point of water removal from the patient in a hemodialysis operation was modelled to identify candidate factors that can be personalized (Pietribiasi *et al.*, 2016; Pietribiasi *et al*., 2018). Modelling has been used to demonstrate mechanisms of solute removal (Werynski & Waniewski, 1995) and uremic toxin removal (Maheshwari *et al.*, 2017) in typical simulated dialysis procedures. Relevant to current focus on reducing immune system storms, the processes underlying cytokine capture have been demonstrated in a recent modelling study (DiLeo *et al.*, 2009). The procedure of dialysis itself has been modelled by Ursino *et al.* (Ursino *et al.*, 2000), which forms a sub-model in the presented study.

### 1.3 Study aims

This work provides an *in silico* system for preliminary assessment of dialysis and therapeutic hypothermia therapies. Specifically, we aimed to implement an open source model of the human circulation coupled to a dialysis circuit; simulate hemodynamic expectations under multiple conditions in model populations; and perform a sensitivity analysis to identify the important parameters in our model.

## 2. Methods

### 2.1. Model components

The composite model consists of a whole body circulation model where baroreflex control is present, which is coupled to a representation of a dialysis machine. A whole body blood circulation model (Heldt *et al.*, 2002; Heldt *et al.*, 2010) with known baroreflex control mechanisms (deBoer *et al.*, 1987; Schuessler *et al.*, 1988; Lin *et al.*, 2013a) was coupled to a virtual dialysis unit (Ursino *et al.*, 2000; Lim *et al.*, 2008). The circulation model was extended to include detailed kidneys. The two kidneys, left and right, were placed in parallel and both received inlet flow from the descending aorta. Further, renal microvasculature was represented within each kidney. Each renal microvasculature bed was represented by a 3 element Windkessel model, and six microvasculature beds in parallel were used to describe the complete microvasculature.

### 2.2. Four model populations

Models populations for baseline (B), therapeutic hypothermia (TH), hemodialysis (HD), and hemodialysis with therapeutic hypothermia (HD&TH) were simulated in this study.

Each model population consisted of 10^4^ instances. To generate model instances, a spectrum of relevant model parameters were perturbed to represent inter-patient variability. Model parameters values were randomly sampled from Gaussian distributions with predefined mean values and a coefficient of variation of 0.25. An implementation of the Box-Muller algorithm (Fernandez & Criado, 1999) ensured Gaussian sampling, while representation through the parameter range was ensured by use of Latin Hypercube Sampling (Calvo *et al.*, 2018). In all simulations, four cases including baseline (B), hemodialysis (HD), therapeutic hypothermia (TH), and hemodialysis under therapeutic hypothermia (HD&TH) were considered. The first of the four model populations were generated under baseline conditions. The HD population was generating by activating clearance of carbonate, urea, and small molecules (potassium, calcium, and sodium) from the blood (Ursino *et al.*, 2000). Further, HD induced increase in heart rate to 90 bpm was implemented. Finally, under HD conditions, ultrafiltration from blood to the dialyzer was activated. Therapeutic hypothermia (TH) treatment was simulated by implementing an experimentally observed Q10 scaling of all model resistances (Konstas *et al.*, 2007). Whereas physiological temperature was taken to be 37.5 C, TH temperature was conservatively assumed to be 35.5 C throughout the body. The following relationship was used to scale all blood vessel diameters with respect to temperature:

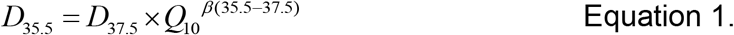

where *D_τ_* represents vessel diameter at temperature *T*, *Q_10_* = 2.961, and *β* = 0.084 (Konstas *et al.*, 2007). The diameters were then used to assign all resistance values (resistance, *R*, is inversely proportional to the fourth power of diameter, *D*). Finally, HD&TH conditions simulated whole body cooling of 2C simultaneous to HD.

### 2.3. Clinically relevant model outputs

Clinically relevant outputs were obtained in all simulation experiments. The outputs were cardiac output, heart rate, systolic and diastolic pressures of the systemic artery (aorta), renal pressures and flows. In addition, time variations of all dynamic quantities were recorded for further processing. Shear as indicator of myogenic stimulant (Arciero *et al.*, 2008) was computed in all blood vessels, and recorded in the aorta (low resistance and high compliance) and kidneys (high resistance). Shear was estimated using the flow (*Q*), diameter (*D*), and blood viscosity (η) as:

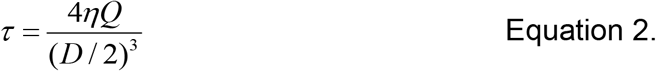

where *τ* is the shear, *η* = 3×10 ^5^ mmHg-s is the constant blood viscosity in all vessels (Heldt *et al.*, 2002), *Q* is flow in the vessel, and *D* is the temperature dependent diameter.

### 2.4. Simulated pathological conditions

In this work, the effects of atrial fibrillation (AF), exercise, and right kidney failure on the efficacy of HD and TH were assessed.

AF was simulated by clamping both atrial elastances to their diastolic values, 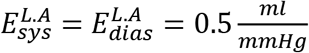 and 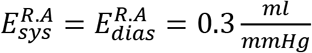. In addition, AF induced irregular heart rate was incorporated into the model simulated as described in the literature (Anselmino *et al.*, 2017).

Exercise was simulated by increasing the ventricular end-systolic elastances and the heart rate, while reducing compartment resistances and large vessel compliances (Anselmino *et al.*, 2017). Specifically, the ventricular systolic elastances were increased by 140%, and the intrinsic heart rate was increased by 28.6%. Simulated exercise also encompassed a reduction of all microvascular resistances by 33%, and a reduction of compliances either by 10% (all aortic segments) or by 32% (all venous segments). (Anselmino *et al.*, 2017).

The consequences of right kidney dysfunction on clinical measurables were simulated. The presence of right kidney dysfunction was programmed as either a large increase of right kidney resistance, or a large simultaneous reduction of aortic and right kidney compliances. Further, right kidney dysfunction was simulated as either homogeneous or inhomogeneous conditions. Under homogenous conditions, all microvascular resistances in the right kidney were either augmented 10 fold, or compliances reduced by 90%, or both resistance and compliance alterations implemented simultaneously. Heterogenous right kidney dysfunction was simulated by manipulating three of the six right kidney microvascular beds’ resistances (increased 10 fold) and compliances (reduced by 90%). In addition, in accordance with experimental information (Amann *et al.*, 1997; Lin *et al.*, 2013b), dysfunction of right kidney was also defined as either a 10 fold increase of the right kidney’s resistance, a 90% reduction of compliance (i.e. increase of stiffness) of the aorta (systemic artery), or both. Results using the latter definition of right kidney dysfunction are presented.

The HD sub-model simulates temporal changes of plasma volume under prescribed ultrafiltration targets. The efficacy of HD was assessed using the fraction change of plasma volume. The efficacy of HD was assessed under AF, exercise, and right kidney dysfunction conditions.

### 2.5. Assessment of parametric sensitivity

Sensitivity of the clinically relevant model outputs (see above) to model mechanism regulating parameters was computed using our implementation of partial ranked correlation coefficients (Marino *et al.*, 2008). The coefficients were used to rank the parameters in descending order of significance, and the most relevant results reported in this study.

### 2.5. Numerical methods

The model was first prototyped using high level programming (MATLAB). It was translated to our C language based differential-algebraic equation solver suitable for efficient large production runs (Altamirano-Diaz *et al.*, 2019). The implicit solver is highly stable and provides high accuracy (*O*(dt^6^)) (Hindmarsh *et al.*, 2005). In all simulations, the maximum time step, dt, was taken to be 0.05 s with tolerances (both, absolute and relative) of 10^-6^ for the adaptive time step solver. Whereas one instance of the model required a maximum of 180 s of serial run-time, a model population consisting of 10^4^ instances was generated within 3 hours using 64 processors. The MATLAB code of the presented model is provided openly in our GitHub (https://github.com/mccsssk2/Frontiers2021) repository.

## 3. Results

### 3.1. Baseline model

The baseline model population’s aortic and kidney pressures as well as flows and shear are illustrated in **Figure 1.** The model’s structure is provided in **Supplementary Materials, Figure S1 (A, B, C, and D).** Representative parameter Gaussian distributions used to generate the baseline population are shown in **Supplementary Materials, Figure S2**.

**Figure 1.**
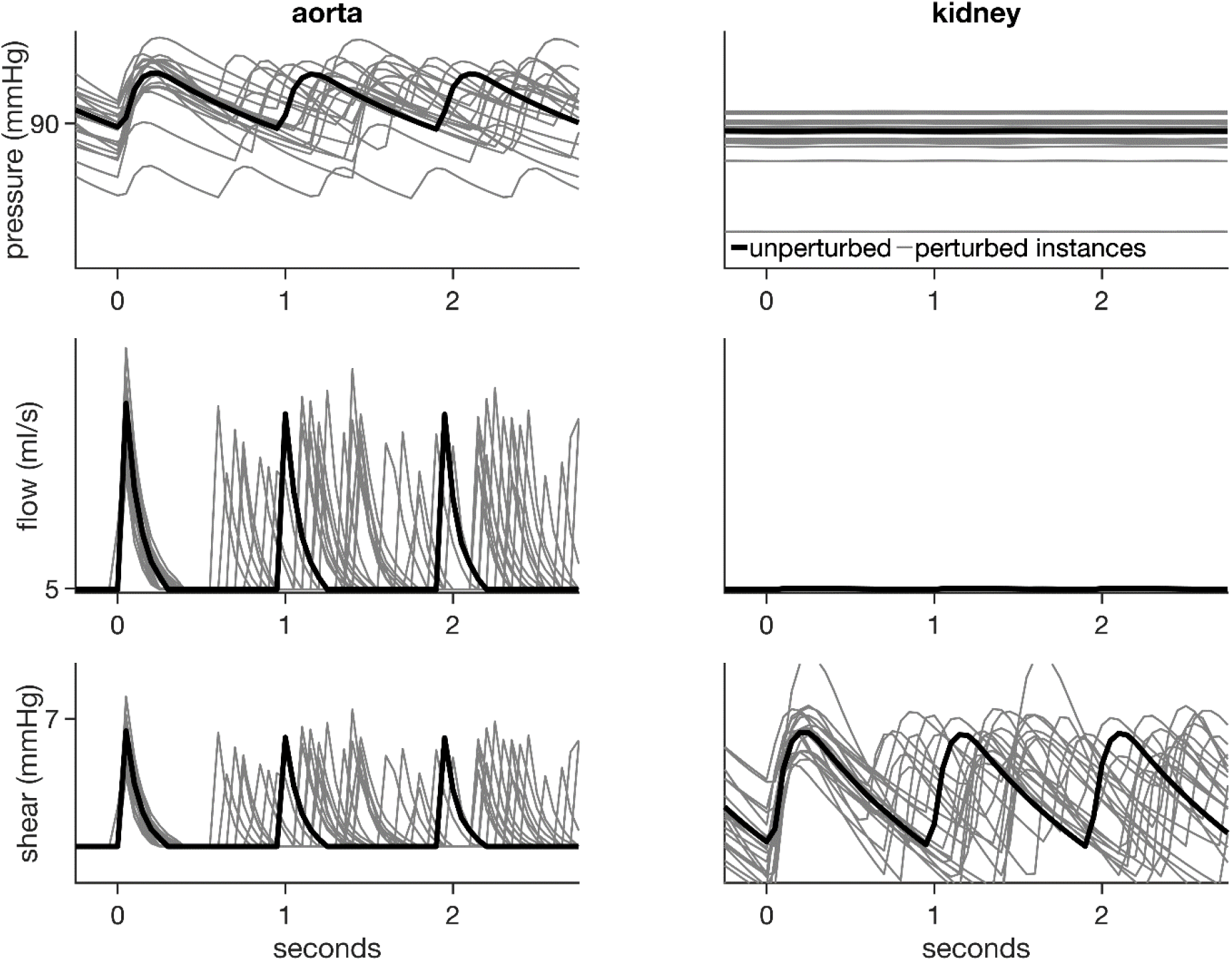
Baseline model. In all panels, black lines show unperturbed data while grey lines show a few (20) representative population instances. Left column shows data for aorta and right column shows data for right kidney. Top row: Pressure profiles. Second row: Blood flows. Bottom row: Shear.

The model systemic artery’s (aorta’s) pressure has a diastolic value of 80 mmHg and a systolic value of 120 mmHg (**Figure 1, top left**). In contrast, the kidney inlet pressure was found to be constant at around 52 mmHg (**Figure 1, top right**). Whereas the aortic blood flow was observed to vary between almost 0 to 1000 ml/s, the kidney’s inlet flow was constant at 4 ml/s (**Figure 1, second row)**. It was also observed that the kidney experienced beat to beat prolonged shear in comparison to the aorta (**Figure 1, bottom row**).

### 3.2 Clinically relevant model outputs

The effects of HD, TH, and simultaneous HD with TH (HD&TH) on clinically relevant measurables (outputs) are illustrated in **Figure 2**. TH led to a reduction in cardiac output (17% under TH, 14% under HD&TH), while HD augmented it by a small amount (3% increase) (**Figure 2, A**). While TH was found to augment heart rate, it was predominantly increased due to HD (**Figure 2, B**, 20% increase under HD). The diastolic and systolic pressures remained largely unaffected by TH, HD, or HD&TH (**Figure 2, C and D**). While shear (Figure 2, E and F) was reduced by TH (15% in the aorta, 6% in the kidney), HD augmented the same (5% in both the aorta and kidney). Under HD&TH conditions, shear was reduced in the aorta by 10% and in the right kidney by 3%. Blood flow distributions in the four model populations (baseline, TH, HD, HD&TH) are illustrated in **Figure S3**, while baroreflex control is shown in **Figure S4**. Whereas all outputs shown in **Figure 2** are derived quantities based on the model’s pressure and flow variables, their spread (represented by the standard deviations) was observed to be affected due to the underlying non-linear dynamical relationships. Overall, the spread of cardiac output and shear under TH and HD reduced, while the spread of heart rate increased. The increase in uncertainty in heart rate was observed to be predominantly due to HD (20% increase of standard deviation, **Figure 2, B, red bar**). All data in sections 3.3 and 3.4 (Figures 3 and 4) are normalized to the respective baseline values of **Figure 2 (Figure 2, gray bars)**.

**Figure 2.**
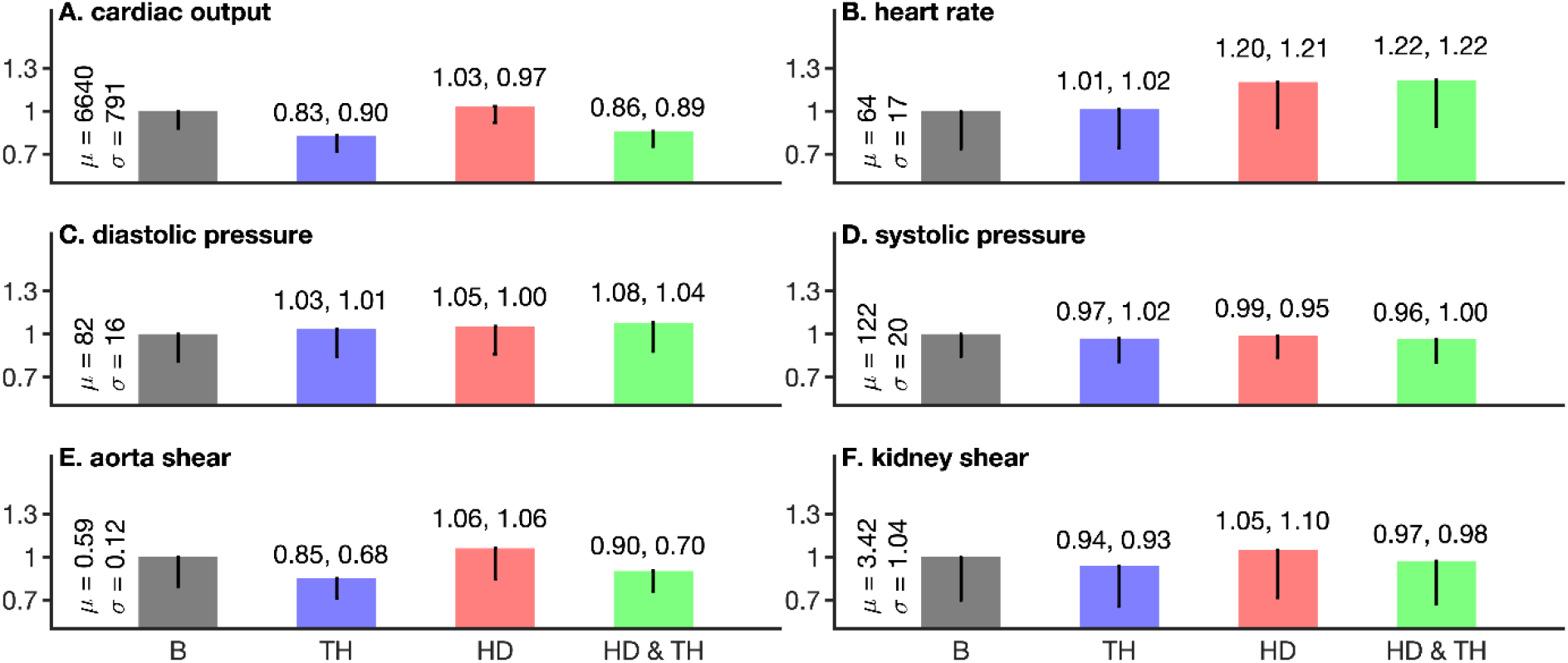
Clinically relevant model outputs in four populations. All bar graphs are normalized to the means of the respective baseline (bars labelled B in all panels) data. Bar colours: Baseline (B, grey), hemodialysis (HD, red), therapeutic hypothermia (TH, blue), and simultaneous hemodialysis with therapeutic hypothermia (HD & TH, green). A: Cardiac output. B: Heart rate. C: Systolic blood pressure. D: Diastolic blood pressure. E: Aortic shear. F: Right kidney shear.

**Figure 3.**
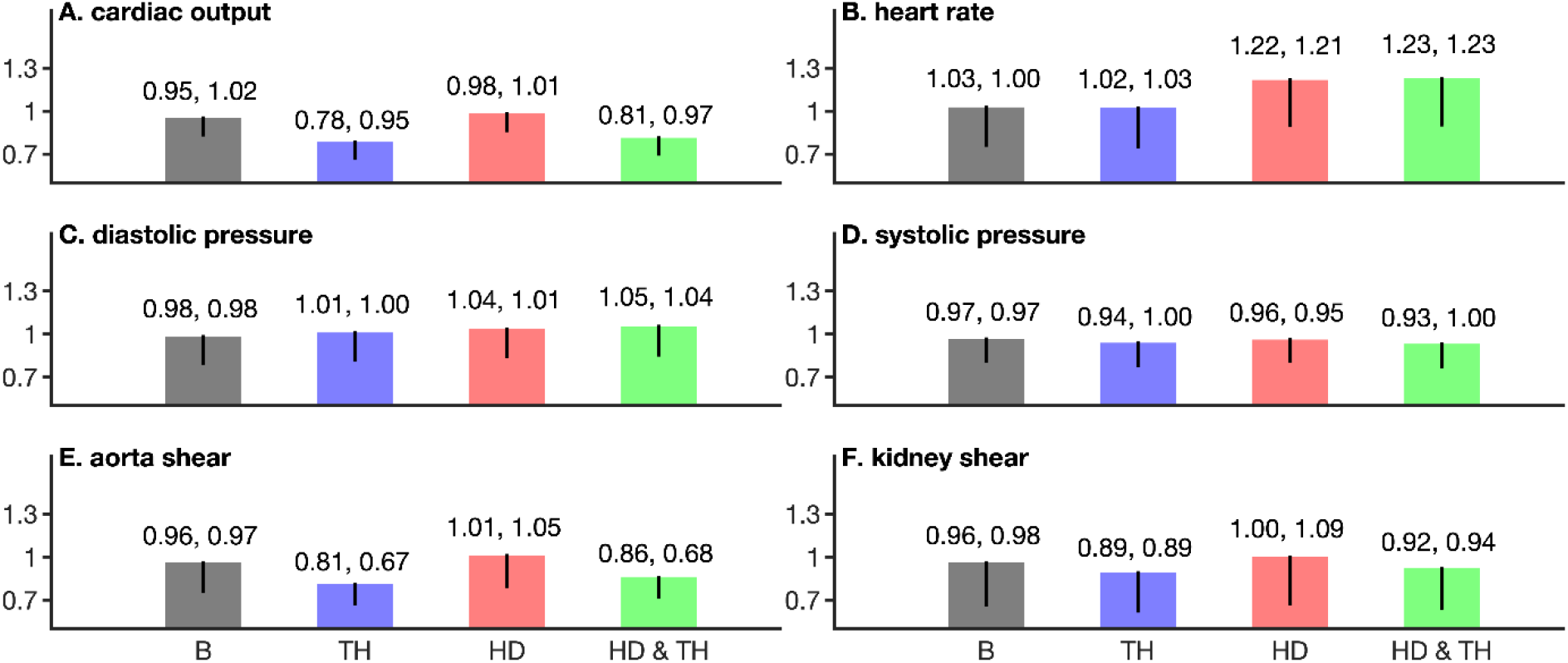
Simulated atrial fibrillation (Anselmino *et al.*, 2017) under baseline (B, gray), therapeutic (TH, blue), dialysis (HD, red), and dialysis under TH (HD&TH, green) conditions. All data are normalized to the respective baseline (bars labelled B in all panels) data of **Figure 2**. A: Cardiac output. B: Heart rate. C: Systolic blood pressure. D: Diastolic blood pressure. E: Aortic shear. F: Right kidney shear.

**Figure 4.**
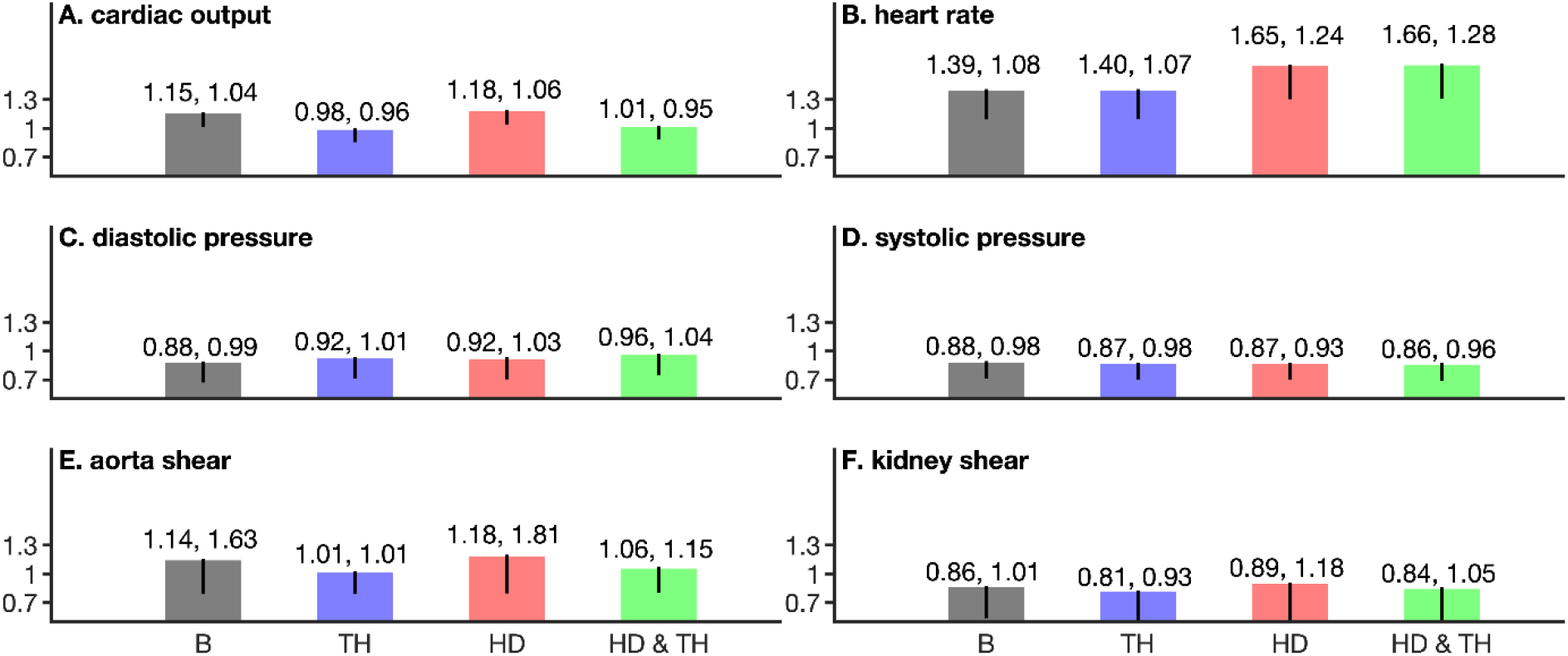
Effects of simulated exercise (Anselmino *et al.*, 2017) under baseline (bars labelled B, gray), therapeutic hypothermia (TH, blue), dialysis (HD, red), and dialysis under TH (HD&TH, green) conditions. All data are normalized to the respective baseline (B) data of **Figure 2** (respective gray bars in **Figure 2**). A: Cardiac output. B: Heart rate. C: Systolic blood pressure. D: Diastolic blood pressure. E: Aortic shear. F: Right kidney shear.

### 3.3. Role of atrial fibrillation

AF conditions, as described in *Section 2.4 Simulated conditions,* were imposed on all four populations. The results, illustrated in **Figure 3,** are presented in comparison to the non-AF case discussed in Sec. 3.2 and shown in **Figure 2**. Under baseline conditions (**Figure 3, gray bars**), AF reduced cardiac output and blood pressure, increased heart rate, and had a heterogeneous effect on shear. In comparison to the non-AF case (**Figure 2**), AF reduced cardiac output in all four populations and the maximal reduction of 20% was under hypothermia (TH) conditions (**Figure 3, A,** blue and green bars). HD has a small reducing effect on cardiac output (**Figure 3, A,** red bar). The changes in spread of cardiac outputs in all AF affected populations was minimal in relation to the non-AF case (**Figure 2**). Under AF, HD predominantly increased the heart rate by over 20% (**Figure 3, B,** red and green bars) in relation to the non-AF baseline case (Figure 2, B). Further, the spread of heart rates was maximally increased by HD under AF conditions (**Figure 3, B,** error-bars of red and green bars). The diastolic and systolic blood pressures remained comparable to the respective baseline cases (**Figure 2, C and D**, gray bars) under AF conditions (**Figure 3, C and D**). AF conditions reduced shear in both the aorta (**Figure 3, E**) and in the kidney (**Figure 3, F**). The reduction was maximal under therapeutic hypothermia (TH) conditions with a maximal reduction of 20% in the aorta and 11% in the kidney. Under HD&TH conditions, the shear augmenting effects of TH were partially abrogated. In all cases (**Figure 3, E and F**), the spread (intra-population standard deviation) of shear data was reduced by TH under AF conditions.

### 3.4. Role of exercise

Exercise conditions were imposed on all four populations, and the model outputs (population means and standard deviations) compared to the non-exercise case of **Figure 2** *(section 3.2).* Under baseline conditions (**Figure 3**, gray bars), exercise increased cardiac output, increased heart rate, reduced blood pressures, and heterogeneously affected the shear. In comparison to the non-exercise case (**Figure 2**), exercise increased cardiac output by over 15% in the normothermic populations (**Figure 4, A,** gray and red bars), and was unaffected by TH (**Figure 4, A,** blue and green bars). The changes in spread of cardiac output in all exercise affected populations was minimal in relation to the non-exercise case (**Figure 2**). Under exercise, HD predominantly increased the heart rate by over 65% (**Figure 4, B,** red and green bars). Heart rate was also increased under baseline-exercise and TH-exercise conditions by 40% (**Figure 4, B**, gray and blue bars). The population wide heterogeneity in heart rates was augmented due to HD (**Figure 4, B**, red and green bars). The diastolic and systolic blood pressures were reduced by 8% in the baseline-exercise case (**Figure 4, C and D,** gray bars). Both TH and HD conditions promoted higher diastolic but not systolic pressures (**Figure 4**, panels C and D). The standard deviations of the population pressures remained unaffected in relation to the non-exercise case of **Figure 2**. Exercise conditions augmented shear in the aorta by 14% (Figure 4, E, gray bar) and reduced it in the kidney by 14% (**Figure 4, F**, gray bar). Application of TH reduced shear in both the aorta (large vessel) and in the kidney (microvasculature) (**Figure 4, E and F,** blue and green bars). HD was seen to be a shear augmenting factor under exercise. The effect of exercise in intra-population heterogeneity (standard deviation) was different among the large aorta and the smaller kidney vessels. In the aorta, the heterogeneity was augmented under baseline-exercise and HD conditions, while the same was reduced in the kidney. TH reduced the shear heterogeneity.

### 3.5. Role of one homogenous kidney dysfunction

Homogenous right kidney dysfunction was simulated by either augmenting all right renal microvascular resistances ten fold, or by reducing aortic compliance by 90% (**Figure 5**). Under non-failing right kidney conditions (**Figure 5, Ai and Aii**), TH reduced inlet flow by 18%. In contrast, HD augmented renal flow by 3% in each kidney. Under TH&HD conditions, the renal flow was reduced by 14% (**Figure 5**, Ai and Aii, green bars) as compared to baseline (**Figure 5, Ai**, gray bar). TH reduced the intra-population heterogeneity of renal flow (**Figure 5, Ai and Aii**), while it was increased by HD.

**Figure 5.**
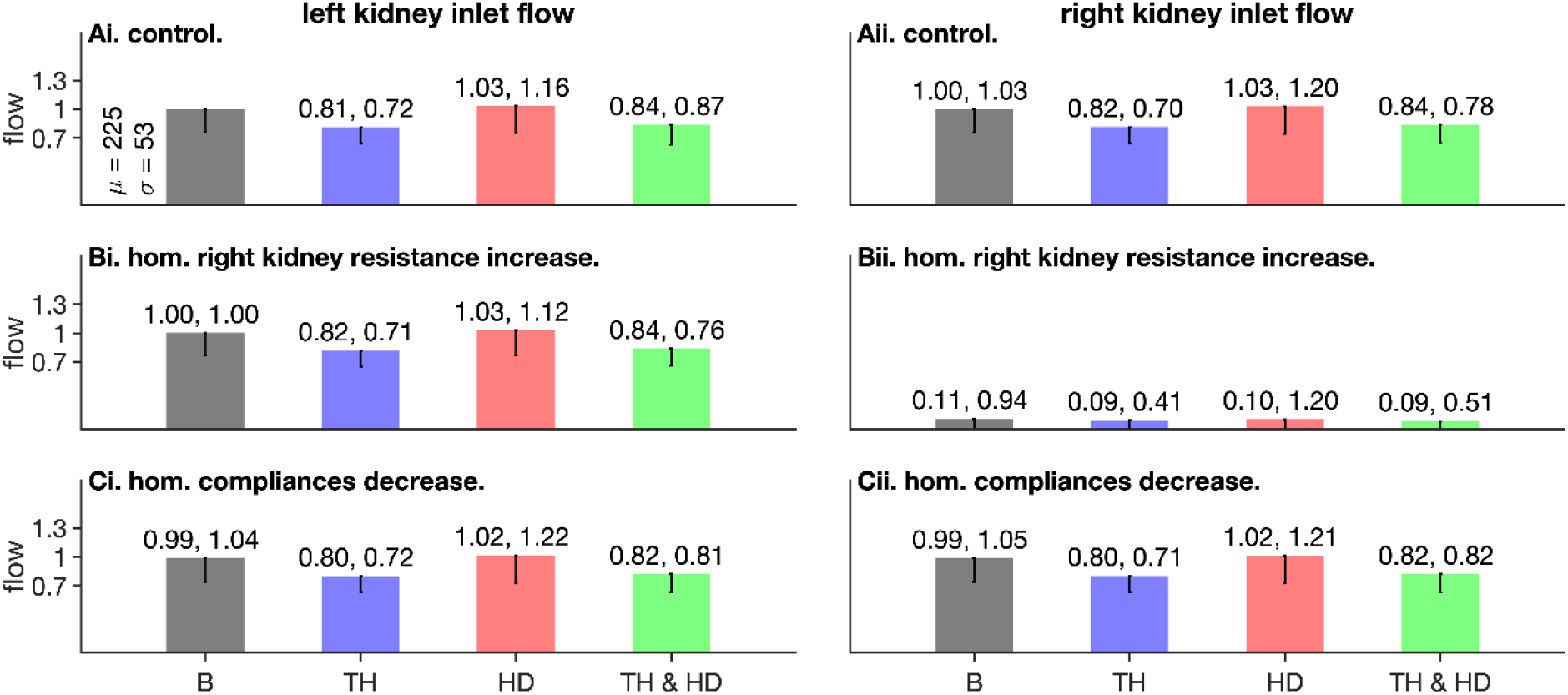
Effects of homogenous right kidney failure on renal inlet flow. Left column shows data for left kidney, right column shows data for right kidney. Top row shows data for control (absence of right kidney failure), middle row shows data when right kidney microvascular resistance was increased 10 fold, and bottom row shows data when right kidney microvascular compliance was reduced (i.e. stiffness was increased) by 90%.

When all microvascular resistances in the right kidney were increased by ten fold, the flow going to the right kidney virtually vanished (**Figure 5, Bii**) and that to the left kidney increased (**Figure 5, Bi**). The effects of TH, HD, and TH&HD on left kidney flow under right renal failure (increased resistances) are shown in **Figure 5, Bi**. In addition, TH reduced intra-population heterogeneity in both the left and right kidneys (reduced standard deviations), and it was augmented by HD.

Reduction of aortic and right kidney compliances did not affect inlet flow to either kidneys (**Figure 5, Ci and Cii**, gray bars). Under TH conditions, both kidneys experienced reduced flow, while HD increased the flow by a small amount. TH&HD conditions (Figure 5, Ci and Ci, green bars) generated an 18% reduction of flow to both kidneys. While HD increased intra-population heterogeneity in flow by 22% due to increased stiffness, TH was found to reduce it by almost 30%. Under TH&HD conditions, the heterogeneity was seen to be reduced in comparison to the respective control value of Figure 5, Ai, gray bar.

### 3.6. Heterogeneous right kidney dysfunction

Populations where half (3 of the six renal microvascular beds in the right kidney) of the right renal microvasculature was increased by 10 fold or compliances reduced by 90% to represent heterogenous right kidney dysfunction. Similar to the homogeneous case (*sec. 3.5*, above), TH reduced renal inlet flow, while HD augmented the same. Heterogeneous right renal dysfunction represented by resistance increase was unable to affect renal inlet flows as much as the corresponding homogeneous case (**Figure 5, Bi and Bii**). Altered compliances had a marginal effect on renal inlet (**Figure 6, Ci and Cii**).

**Figure 6.**
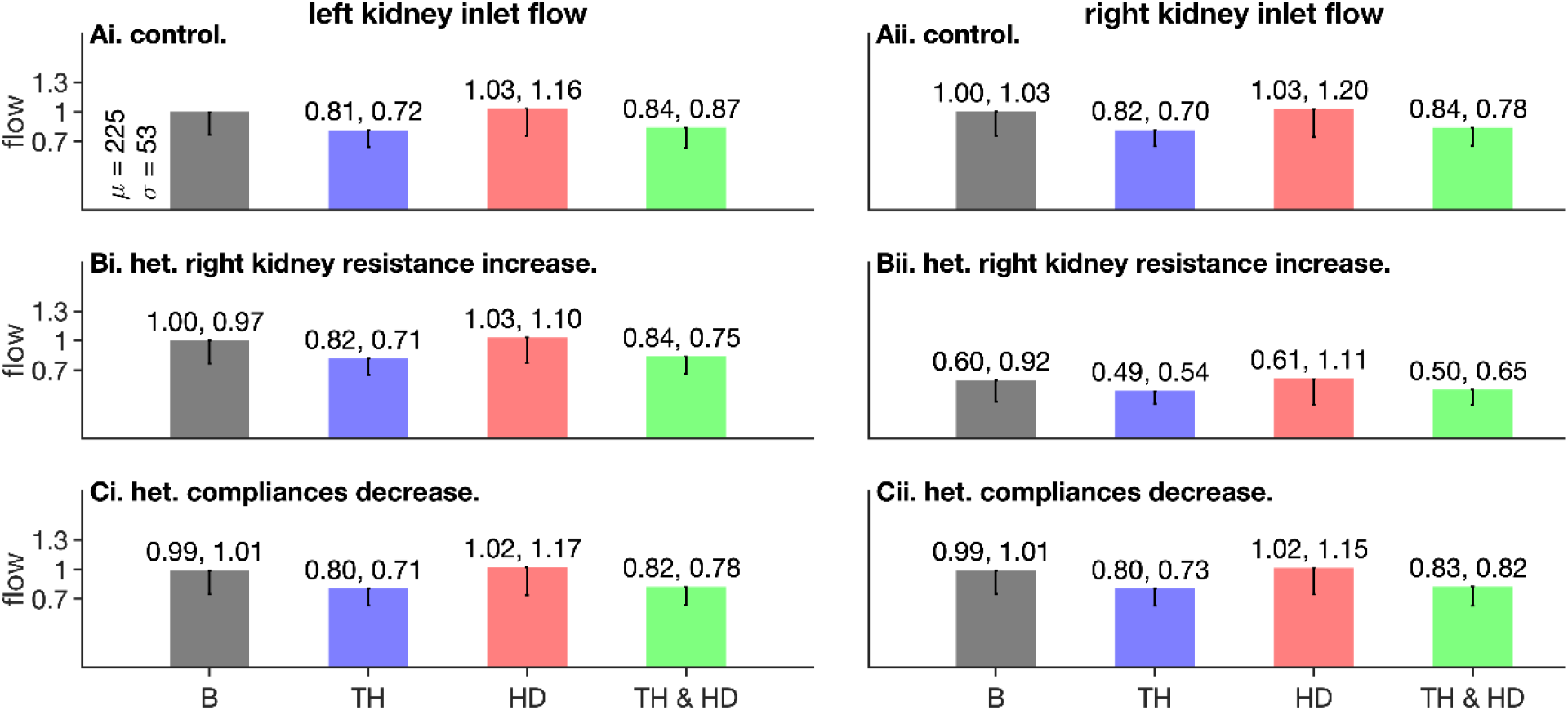
Effects of heterogeneous right kidney failure on renal inlet flow. Left column shows data for left kidney, right column shows data for right kidney. Top row shows data for control (absence of right kidney failure), middle row shows data when half (3 of 6) right kidney microvascular resistances were increased 10 fold, and bottom row shows data when half (3 of 6) right kidney microvascular compliances was reduced (i.e. stiffness was increased) by 90%.

The more significant role of microvascular resistance over compliance is illustrated in **Figure 7**. At each value of compliance and resistance, the six flows in each kidney were noted, and their heterogeneity representing standard deviation used to construct the result (**Figure 7**). As is evident, resistance readily promoted heterogeneity in the affected kidney (right kidney), while compliance plays a marginal role in the presented model.

**Figure 7.**
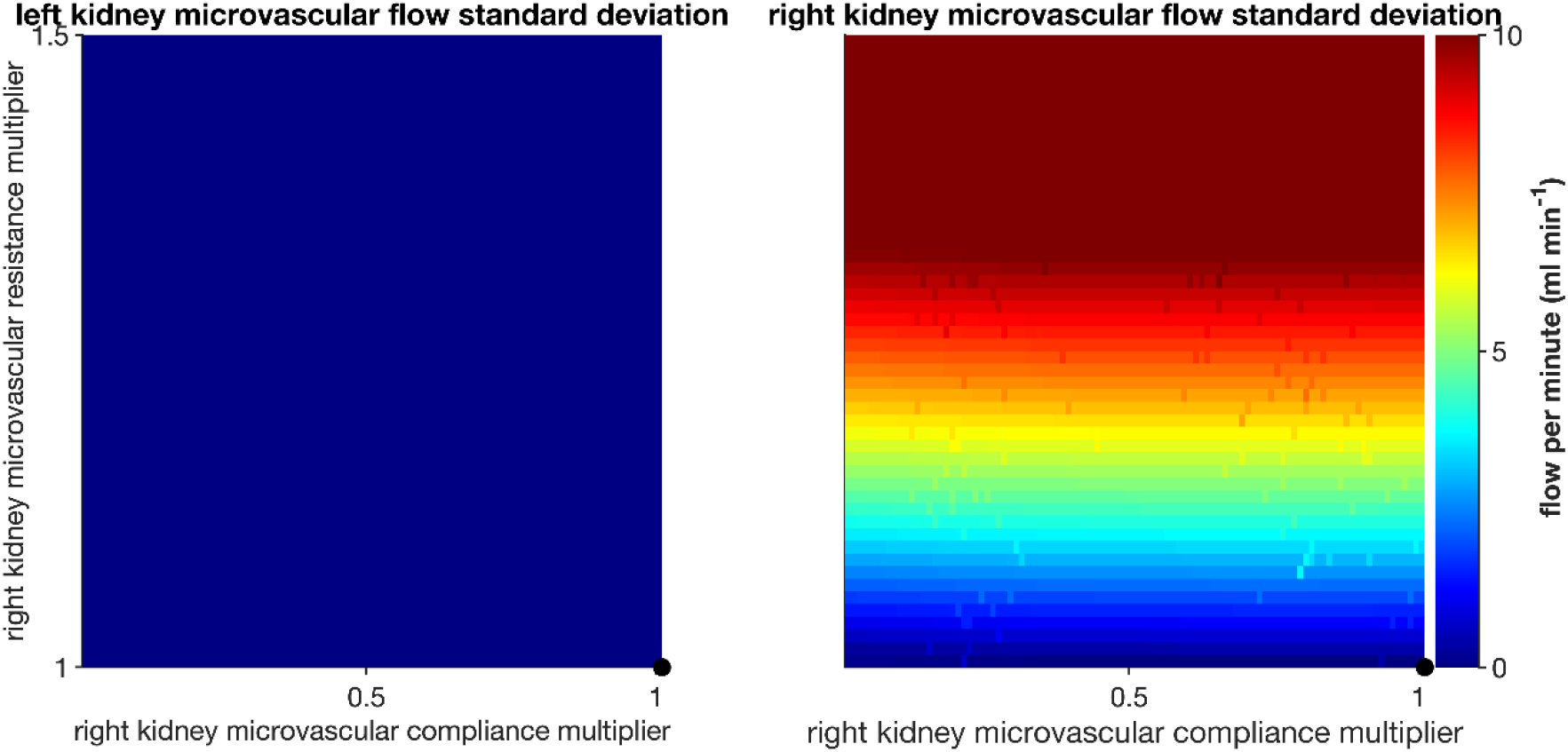
Perfusion heterogeneity in the left and right kidneys due to altered microvascular compliances and resistances. Color coding shows the standard deviation of the microvascular flow for each value of compliance and resistance.

### 3.7. Effectiveness of volume removal under diseased conditions

Plasma volume removal under HD was assessed by the relative change of plasma volume before and after a HD session. Under normothermia and standard target ultrafiltrate (**Figure 8, Ai**), atrial fibrillation, exercise, as well as right renal failure reduced HD efficacy, as reflected in the reduced changes to plasma volume. The effect of exercise was maximal where HD removed 40% less volume as compared to the baseline case (**Figure 8, Ai,** purple and gray bars respectively). TH did not affect the plasma volume removal (**Figure 8, Aii**). Under low target UF (**Figure 8, Bi and Bii**), the alteration of plasma volume was higher and under high target UF (**Figure 8, Ci and Cii**) the alteration of plasma volume was lower. TH was not seen to have any role in plasma volume removal efficacy in the presented model.

**Figure 8.**
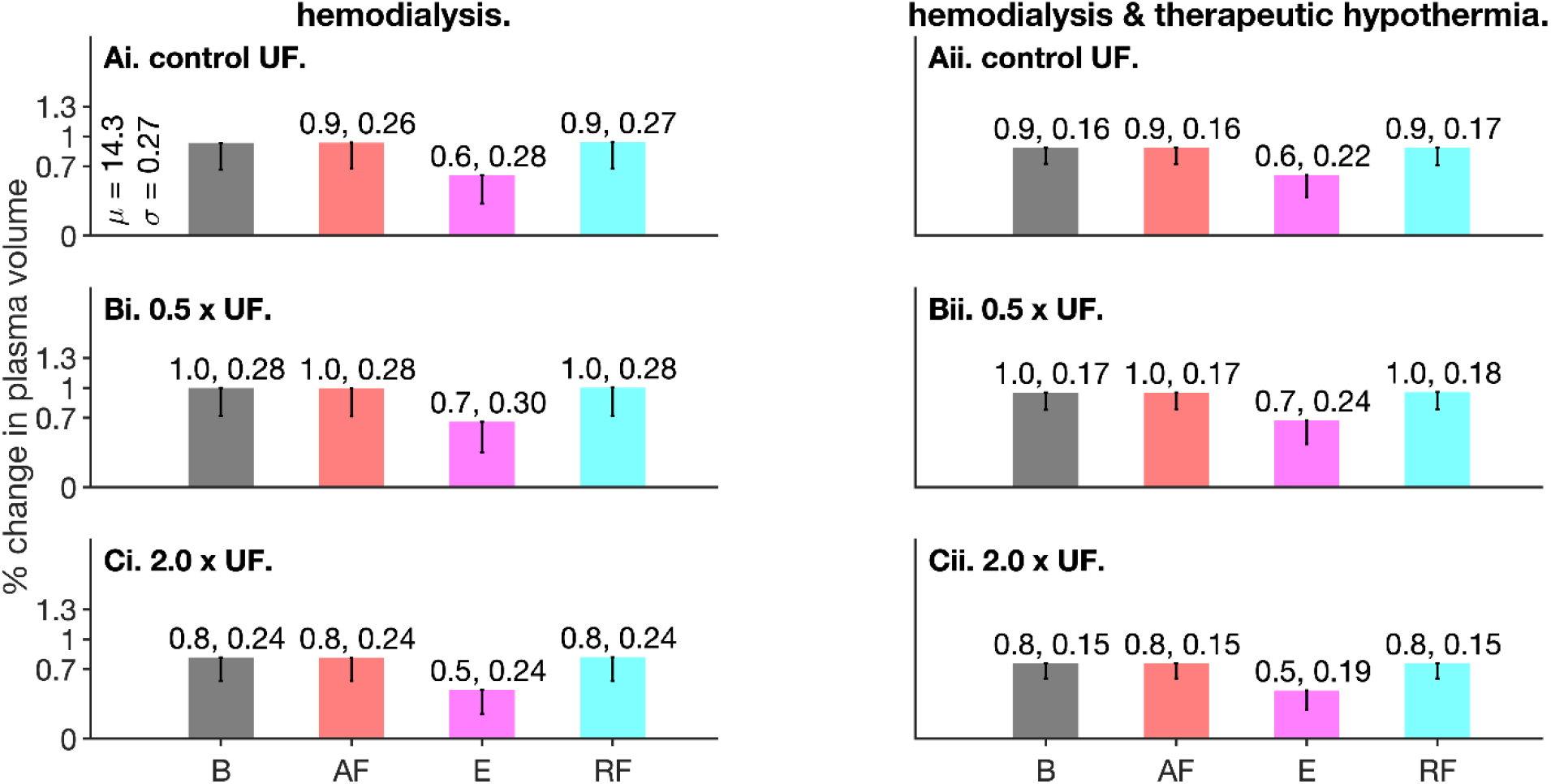
Effectiveness of volume removal due to HD under control and disease conditions and target ultrafiltrate (UF). In all panels, gray (bars labelled B) represent the baseline population, red represents the AF population, purple represents the exercise (bars labelled E) population, and cyan represents the right kidney failure (RF) population. All bars and standard deviations are normalized to the mean value of the baseline population (gray bar’s mean value) in panel Ai.

In comparison to control (**Figure 8, Ai**, gray bar), all conditions reduced the intra-population heterogeneity at all target UF values. Further, TH further reduced the heterogeneity of HD efficacy (**Figure 8**, right column).

### 3.8. Most significant model parameters

The results of PRCC based sensitivity analysis are illustrated in **Figure 9**. Cardiac output (**Figure 9, A**) is positively modulated by the intrinsic heart rate and systolic elastance of the right ventricle under all four conditions (baseline, HD, TH, and HD&TH), while it is negatively modulated by the diastolic right ventricle elastance, upper body resistance, and the baroreflex mechanism parameter, G. Additionally, HD, TH, as well as TH&HD markedly alter the sensitivity of cardiac output to the right ventricle elastance as well as the baroreflex control mechanism. The heart rate (**Figure 9, B**) is maximally modulated by the intrinsic heart rate and the baroreflex mechanism, while resistances (splanchnic and lower body) and right ventricle diastolic elastance play a secondary but significant role. While heart rate depends positively on the intrinsic heart rate and diastolic right ventricle elastance, it is modulated negatively by the baroreflex and resistance mechanisms. The systemic artery’s systolic (**Figure 9, C**) as well as diastolic (**Figure 9, D**) pressures are maximally modulated by the baroreflex. Resistances and elastances play an important role in modulating both. Further, the systemic artery’s compliance (aortic compliance) negatively modulates the systolic pressure while it positively modulates diastolic pressure (**Figure 9, C and D**). Blood vessel shear was seen to be modulated by intrinsic heart rate, baroreflex, elastances, and microvascular resistances. The ranked PRCC sensitivities to shear are shown in **Figure 9, E and F**.

**Figure 9.**
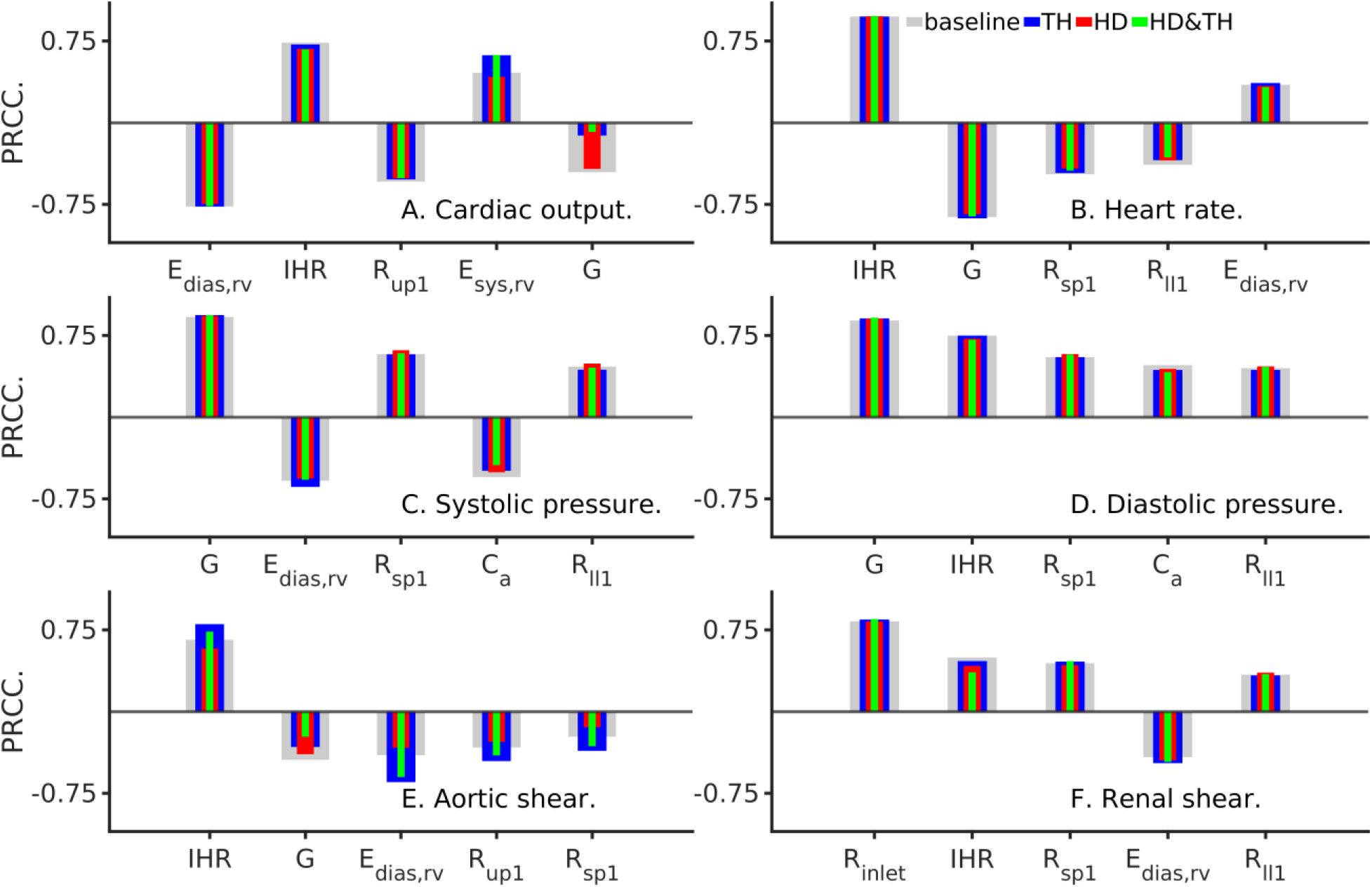
Sensitivity of clinically relevant model outputs to parameters. In all panels, E_dias,rv_ represents diastolic right ventricle elastance; E_sys,rv_ represents systolic right ventricle elastance; IHR represents intrinsic heart rate; R_up1_ represents upper body resistance; G represents baroreflex regulatory parameter; R_sp1_ represents splanchnic resistance; R_||1_ represents legs resistance; C_a_ represents aorta compliance; Rinlet represents kidney inlet resistance.

## 4. Conclusions and Discussion

A model that combined whole body circulation with detailed kidneys representation, along with baroreflex control and dialysis unit was developed. While the present model derives from previous developments by others (Coli *et al.*, 1998; Heldt, 2006; Lim *et al.*, 2008), this work used it in assessing a therapy (therapeutic hypothermia) pertinent to our clinical research. As seen in **Figure 1**, our implementation is capable of qualitatively reproducing flow-pressure waveforms (Olufsen *et al.*, 2000), in addition to providing difficult to measure information such as shear as well as population wide expectation.

In line with our clinical observations, the model suggests that HD augments cardiac output, heart rate, as well as shear (**Figure 2**). While HD induced augmentation of heart rates and shears may be undesirable side effects (Severi *et al.*, 2001), TH appears to impart benefit by reducing heart rates in agreement with clinical finding (Selby *et al.*, 2006), and normalize vascular shear.

Atrial fibrillation (AF) is a prevalent condition among critically ill patients such as those with renal failure. The presented model suggests that AF potentially exacerbates the HD induced augmentation of heart rate (**Figure 3, B**) and lowers blood pressure (**Figure 3, C and D**). Therapeutic hypothermia neither reduced the HD induced heart rate increase, nor did it reverse the HD induced lowered blood pressure. However, the model does suggest that shear in blood vessels, due to its lower intra-population heterogeneity, is strongly linked to underlying AF. The simulated AF had a marginal effect on model behaviour, an observation that is aligned with the findings of Anselmino et al. (Anselmino *et al.*, 2017). The benefit of TH under AF conditions appears to be reduction of both shear and intra-population heterogeneity.

The modelled exercise conditions (**Figure 4**) increased cardiac outputs but beneficially lowered systolic blood pressure. TH imparts further benefit under exercise by primarily in reducing large vessel (aortic) as well as small vessel (renal) shear, which may provide hemodynamic stability.

Within the confines of the model, renal resistance appears to affect renal flow more than renal compliance (**Figures 5 and 6**). The present findings are aligned with our previous findings (Altamirano-Diaz *et al.*, 2019) where large vessel compliance and small vessel resistance was found to affect overall whole body hemodynamics. Our recent clinical observations in renal failure patients show that dialysis increased blood flow heterogeneity in visceral organs, while therapeutic hypothermia may reduce it. **Figures 5** and **6** show that the cause may be found within the population wide heterogeneity measure of standard deviation, or in the absolute measurement of microvascular flow heterogeneity in the organ (**Figure 7**).

The effectiveness of dialysis was assessed using relative changes of plasma volume (**Figure 8**). As a simple definition, a larger fraction of plasma volume removed from the body by the dialyzer was defined as more effective HD. TH reduced the amount of plasma volume removed. In addition, pathological alterations of the hemodynamics such as atrial fibrillation and renal failure were also seen to reduce the amount of plasma volume to be removed by dialysis. Surprisingly, exercise caused the minimal amount of plasma volume to be removed.

Sensitivity analysis, using partial ranked correlation coefficients, permitted ranking of model parameters. Among the large number of model parameters, a handful emerged to be the most relevant (**Figure 9**). Identifying the parametric regulation of outputs will permit future development of the model mechanisms. As expected, the baroreflex control (represented by parameter G) regulates multiple model outputs of interest. Furthermore, the sensitivity of the model to baroreflex control appears to be reduced significantly by TH. The sensitivity of blood vessel shear also appears to be affected by the right ventricle elastance, as well as peripheral resistances (splanchnic and upper body).

## 5. Limitations

The present model is primed for further development. Implementation of detailed organ level vasculature representations of cerebral (Schollenberger *et al.*, 2020), hepatic (Debbaut *et al.*, 2014), pulmonary-respiratory (Hasler *et al.*, 2019), cario-pulmonary (Albanese *et al.*, 2016), and other will better complement future clinical-modelling studies. Future inclusion of myogenic hemodynamic regulation (Carlson & Secomb, 2005; Arciero *et al.*, 2008) will extend the spectrum of phenomena explored (Arciero & Secomb, 2012) and further refine the presented results. In addition to baroreflex control, vascular tone is known to be significantly regulated by renal function. The inclusion of biochemical mechanisms of gas exchange and metabolism will extend the model to permit drug testing, in addition to the presently tested physical therapy of TH. Complementary to phenomenological hemodynamics, a significant proportion of mechanisms at tissue and cellular levels are expected to be valuable in our future modelling-clinical-imaging approach. We are currently constructing multiscale cell to vasculature models driven by current understanding (e.g. see (Jackson, 2000)) that will assist the dissection of pathophysiological disease mechanisms and pharmacological targets. The mechanisms and pathways of widespread sepsis infections during HD could be further illuminated using the multi-scale models (Calmelet *et al.*, 2014).

## Data Availability

The model source code is made openly available via GitHub. All other data will be shared unreservedly upon request.

## Acknowledgements

This work was supported by Canada Canarie Inc. (project: RS3-111), partly by Canadian Heart and Stroke Foundation grant (G-20-0028717, PI: CWM), and partly by Canadian NSERC grant (R4081A03, PI: Prof. Daniel Goldman). This work is part of OS’s research based undergraduate training in Western University. SRK also thanks Lawson Health Research Institute, Western University, MITACS Globalink (Canada), and Compute Canada for compute, labour, and other resources. The authors thank Dr. Kapiraj Chandrabalan for editing support.

## Author contributions

Study design: JJJ, SRK, DG, and CWM. Model development: SRK, JJJ, and TJH. Simulations: JJJ and SRK. Data analysis and figures: JJJ. Draft writing: JJJ, CS and SRK. Draft completion: SRK. Final manuscript: SRK, DG, and CWM. Funding secured by: CWM, DG, and SRK.

