## Supplement for "Using a Human Circulation Mathematical Model to Simulate the Effects of Hemodialysis and Therapeutic Hypothermia"

*Running title:* Human circulation and dialysis 0D model.

*Keywords:* Lumped parameter model, human circulation, dialysis, ordinary differential equations, sensitivity analysis.

[*]Correspondence:

Drs. CW McIntyre and Sanjay R. Kharche.

Room ELL126C, Kidney Clinical Research Unit, Lawson Health Research Institute, Zone E, Victoria Hospital, 800 Commissioners Road East, London, Ontario, Canada N6A 5W9.

.

**A. Supplementary Sections.**

*Section S1. Model components and equations.*

Our model consists of three sub-models that are coupled to each other. The body circulation model (Heldt *et al.*, 2002), dialyzer model (Coli *et al.*, 1998; Ursino *et al.*, 2000), and the baroreflex models (deBoer *et al.*, 1987; Lin *et al.*, 2013) are illustrated in Supplementary Figure S1, A, B, C, and D. Unless otherwise stated, model parameters have been adapted from the original literature. Inter-model coupling was implemented as described in the literature (Lim *et al.*, 2008). The body circulation model's renal Windkessel representation was further developed in this study. Specifically, the original model's renal arteriolar and microvascular resistances were distributed into two parallel circuits as shown in Figure S1, B. Further, the capacitances were evenly distributed among the two parallel kidneys. In each kidney, the baseline inlet resistance was assigned a value 5 (mmHg-s/ml), and each of the six microvascular resistances were assigned values 15 (mmHg-s/ml). Concurrently, the baseline inlet capacitance was assigned a value 5 mL/mmHg, and each of the six microvascular capacitances were assigned values 1.2 mL/mmHg.

In the coupled model, any resistance with value over 0.1 (mmHg-s/ml) was assumed to be large. All large resistances were dynamically computed as functions of vessel radius (represented by r) and blood viscosity (represented by eta) as:

$$R = \frac{8\eta l}{\pi r^4}$$

Where R is resistance, eta is blood viscosity, l is length of vessel, and r is radius. The radius was considered to depend on temperature as

$$r_{35.5} = r_{37.5} \times Q_{10}{}^{\beta(35.5-37.5)}$$

where $Q_{10}$ = 2.1 is the a constant. In all small vessels (vessels with resistance more than 0.1 mmHg-s/ml), the shear stress was computed as:

$$\tau = \frac{4\eta Q}{r^3}$$

where Q is the blood flow (ml/s) in the vessel.

In this study, we considered cardiac output (CO), systemic artery systolic and diastolic pressures, and heart rate (beats per minute) as clinically relevant outputs. The cardiac output was computed by summing the left ventricle outflow over one heart beat.

Time varying elastance $E(t)$, as a function of diastolic elastance $E_d$, and systolic elastance $E_s$ of each chamber:

$$E(t) = E_d + \frac{E_s - E_d}{2} \ \alpha(t)$$

where $\alpha(t)$ is the activation function, specific to each chamber:

$$\alpha_{ventricle}(t) = \begin{cases} 1 - \cos\left(\pi \dfrac{t - T_{av}}{T_{s,v}}\right) & , \quad T_{av} < t \leq T_{av} + T_{s,v} \\ 1 + \cos\left(2\pi \dfrac{t - (T_{av} - T_{s,v})}{T_{s,v}}\right) & , \ T_{av} + T_{s,v} < t \leq T_{av} + \dfrac{3}{2} T_{s,v} \\ 0 & \end{cases}$$

$$\alpha_{atrium}(t) = \begin{cases} 1 - \cos\left(\pi \dfrac{t}{T_{s,a}}\right) & , \quad 0 < t \leq T_{s,a} \\ 1 + \cos\left(2\pi \dfrac{t - T_{s,a}}{T_{s,a}}\right) & , \quad T_{s,a} < t \leq \dfrac{3}{2} T_{s,a} \\ 0 & \end{cases}$$

The atrio-ventricular time-delay $T_{av}$ = 0.2 s, along with the ventricular systolic time duration $T_{s,v}$ and atrial systolic time duration $T_{s,a}$, all scale with $\sqrt{T}$ where $T$ is the cardiac cycle duration (in seconds).

**Table S1.** Systolic and diastolic elastance values in the model heart.

| | Atria | | Ventricles | |
|---|---|---|---|---|
| | **Left** | **Right** | **Left** | **Right** |
| $\boldsymbol{E_d}$ $\left[\frac{\boldsymbol{mmHg}}{\boldsymbol{ml}}\right]$ | 0.5 | 0.3 | 0.13 | 0.07 |
| $\boldsymbol{E_s}$ $\left[\frac{\boldsymbol{mmHg}}{\boldsymbol{ml}}\right]$ | 0.6 | 0.74 | 2.5 | 1.3 |
| $\boldsymbol{T_s}$ $[\boldsymbol{seconds}]$ | 0.25 | | 0.37 | |

### C. Supplementary figures.

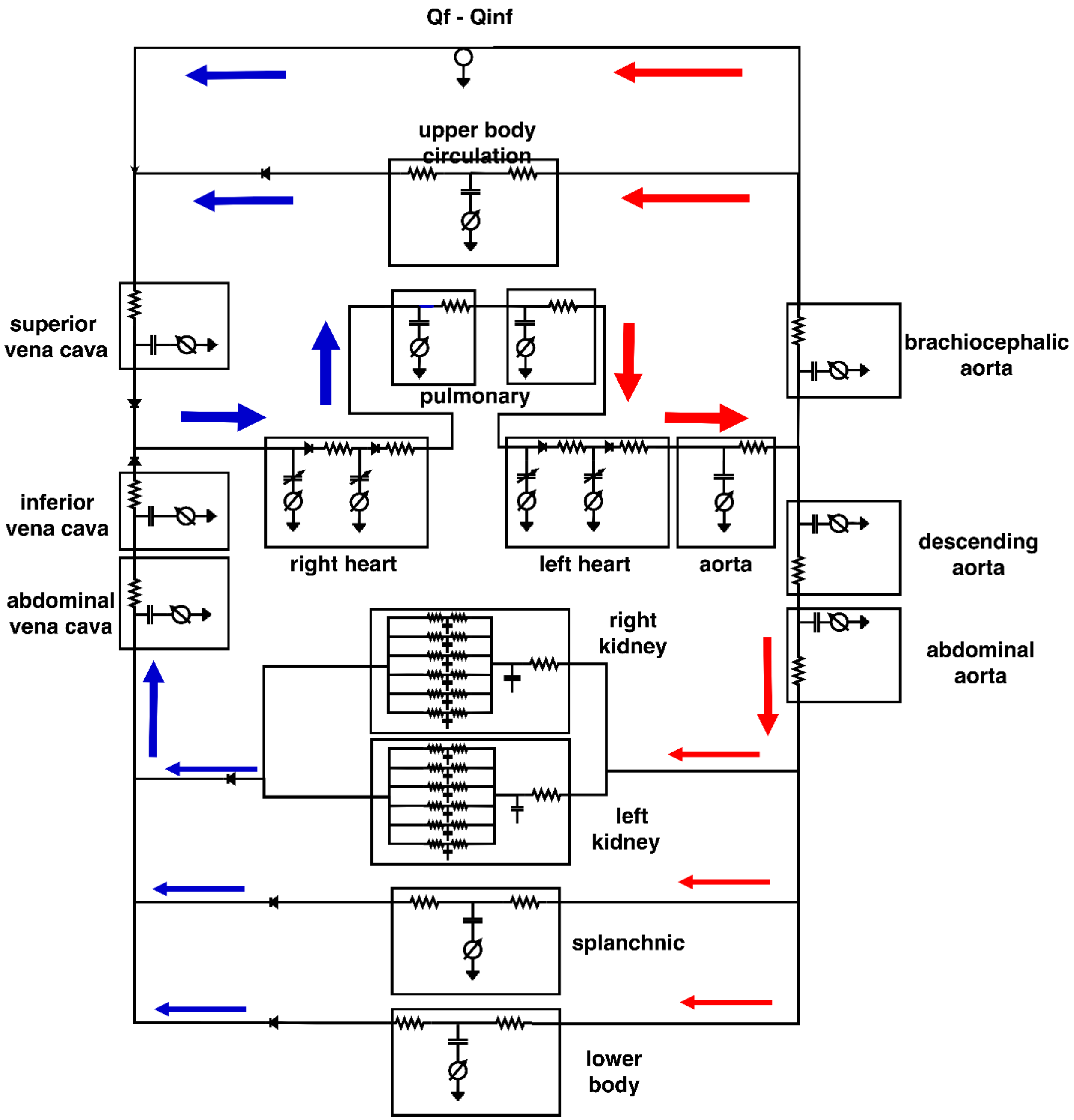


**Figure S1, A.** Schematic of the model’s systemic circulation based on the Heldt model. Arrows represent direction of blood flow in arteries (red) and veins (blue).

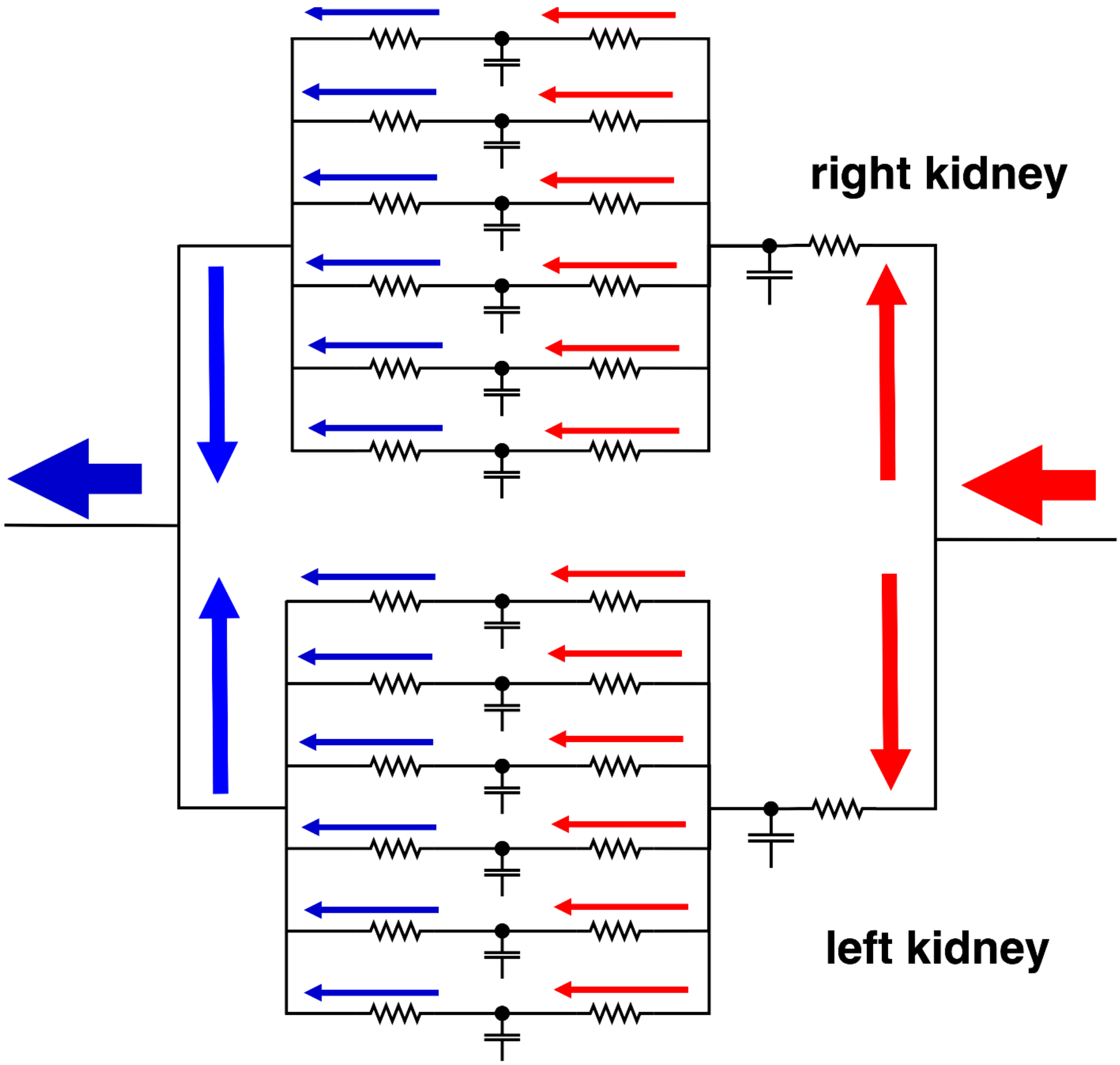


**Figure S1, B.** Schematic diagram of new detailed right (top) and left (bottom) kidneys' complex. Each kidney consists of six microvascular components, implemented as three element Windkessel models. Each component consists of two microvascular beds that is assigned two resistance values and one compliance value.

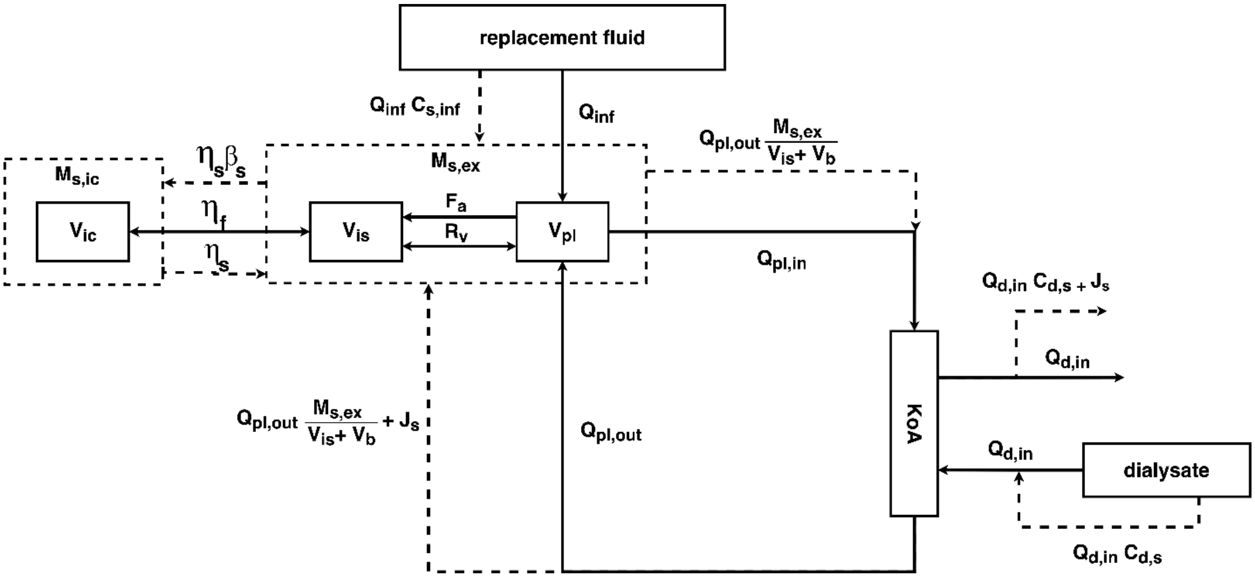


**Figure S1, C.** Schematic of the dialysis unit adapted from the literature (Coli *et al.*, 1998; Coli *et al.*, 2000; Ursino *et al.*, 2000).

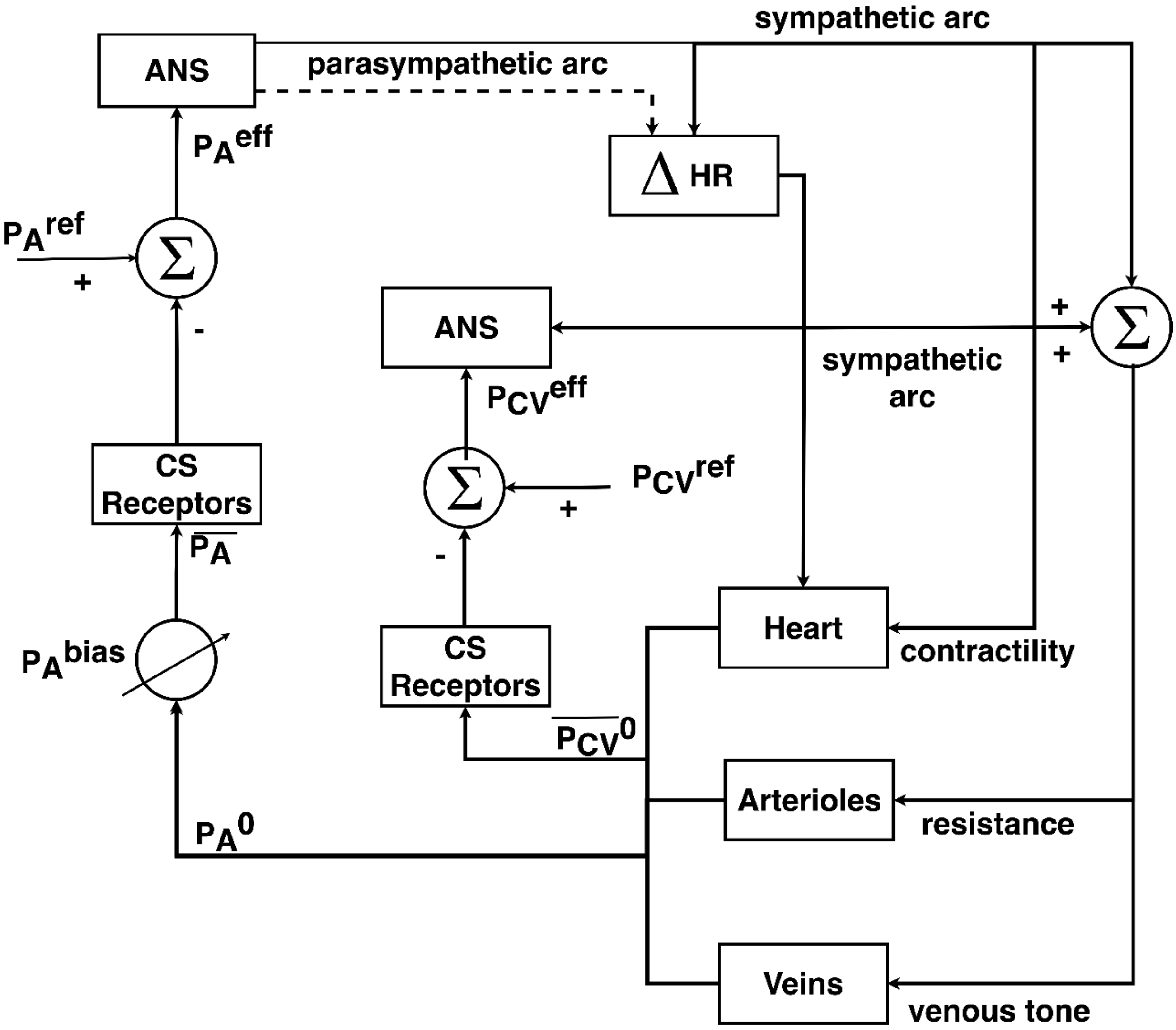


**Figure S1, D.** Schematic of the baroreflex component adapted from Lin et al. (Lin *et al.*, 2012)

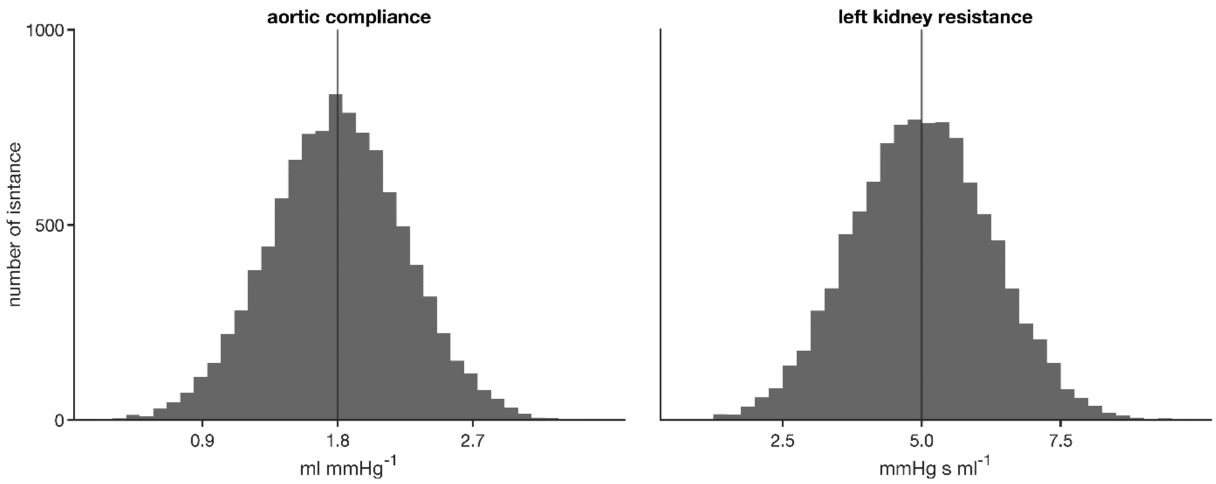


**Figure S2.** Representative parameter Gaussian distributions in the baseline model population (see **Figure 1**, main manuscript) where mean values are taken from the unperturbed model, and a coefficient of variation of 0.25 is used to generate the population. Left panel shows distribution of aortic compliance, and right panel shows distribution of left kidney inlet resistance.

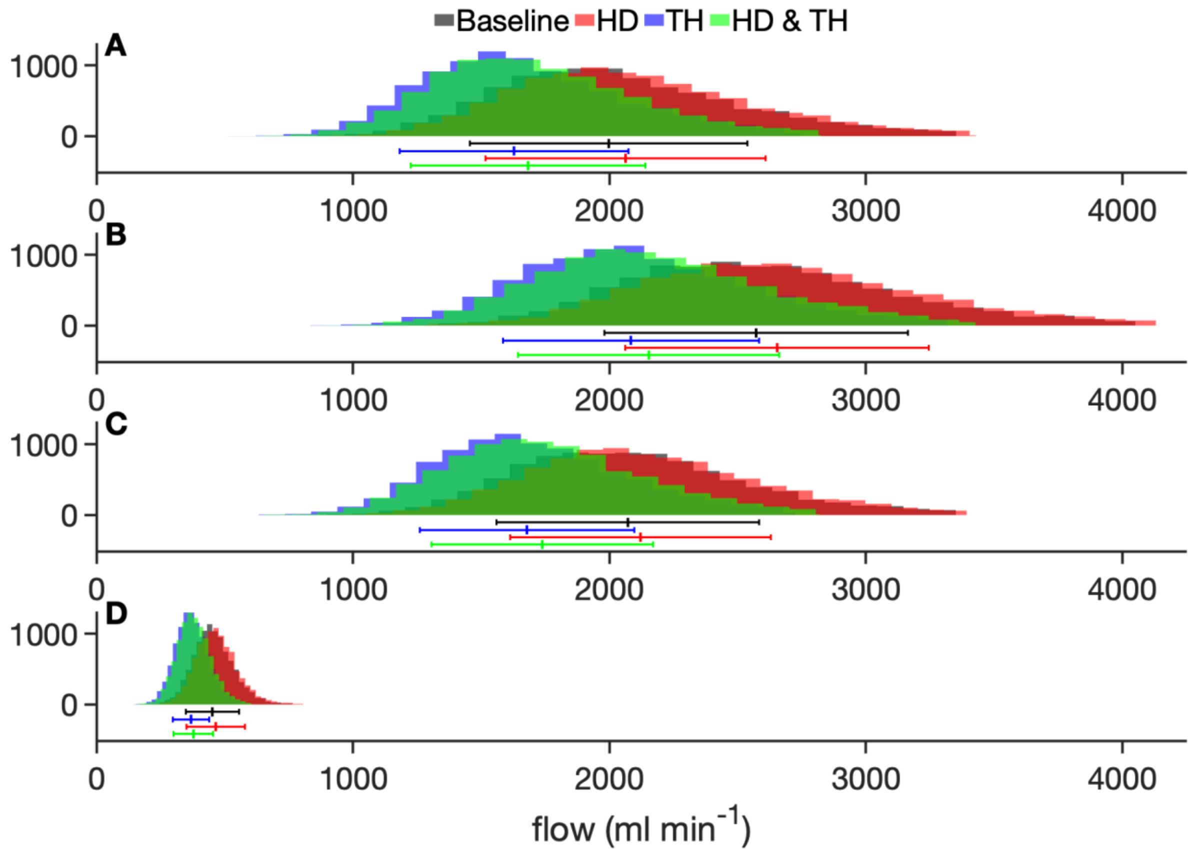


**Figure S3.** Distribution of flows in model organs under baseline (grey), hemodialysis (red), therapeutic hypothermia (blue), and simultaneous hemodialysis with therapeutic hypothermia (green). Figures 1 and 2 in main manuscript. Top row: upper body; second row: splanchnic; third row: lower body; bottom row: both kidneys.

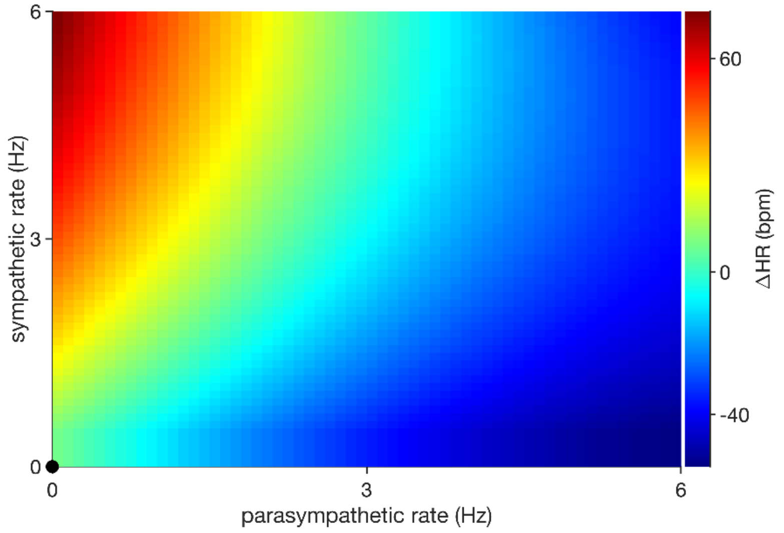


**Figure S4.** Baroreflex control of heart rate by the parasympathetic and sympathetic tones.

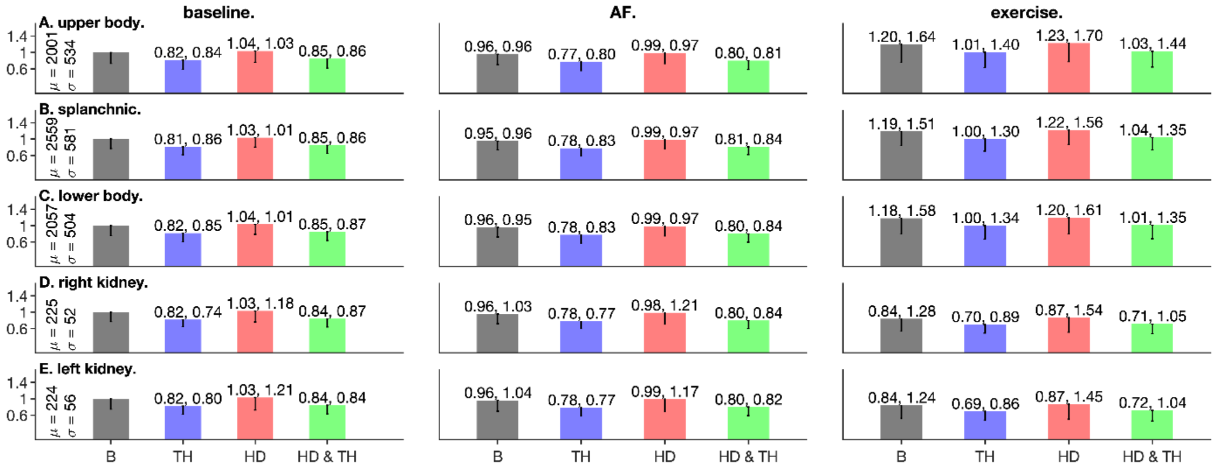


**Figure S5.** Flows under sedentary (left column), AF (middle column), and exercise (right column) conditions. All data are normalized to the baseline values of the left panel.

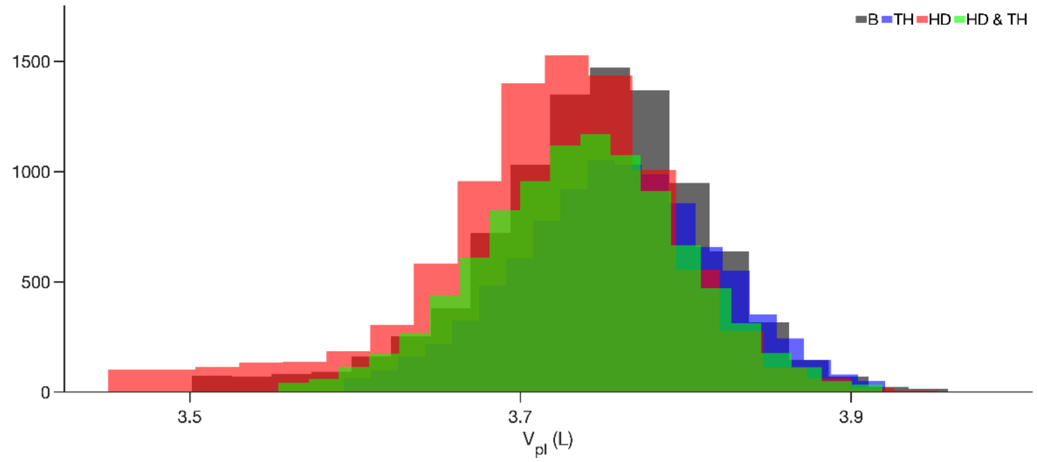


**Figure S6.** Population distributions of intracellular volume (left panel), interstitial volume (middle panel), and plasma volume (right panel). See **Figures 1 and 2** and related text for details.

### References.

Coli L, Ursino M, Dalmastri V, Volpe F, La Manna G, Avanzolini G, Stefoni S & Bonomini V. (1998). A simple mathematical model applied to selection of the sodium profile during profiled haemodialysis. *Nephrol Dial Transplant* **13,** 404-416.

Coli L, Ursino M, De Pascalis A, Brighenti C, Dalmastri V, La Manna G, Isola E, Cianciolo G, Patrono D, Boni P & Stefoni S. (2000). Evaluation of intradialytic solute and fluid kinetics. Setting Up a predictive mathematical model. *Blood Purif* **18,** 37-49.

deBoer RW, Karemaker JM & Strackee J. (1987). Hemodynamic fluctuations and baroreflex sensitivity in humans: a beat-to-beat model. *Am J Physiol* **253,** H680-689.

Heldt T, Shim EB, Kamm RD & Mark RG. (2002). Computational modeling of cardiovascular response to orthostatic stress. *J Appl Physiol (1985)* **92,** 1239-1254.

Lim KM, Choi SW, Min BG & Shim EB. (2008). Numerical Simulation of the Effect of Sodium Profile on Cardiovascular Response to Hemodialysis. *Yonsei medical journal* **49,** 581-591.

Lin CL, Tawhai MH & Hoffman EA. (2013). Multiscale image-based modeling and simulation of gas flow and particle transport in the human lungs. *Wiley interdisciplinary reviews Systems biology and medicine* **5,** 643-655.

Lin J, Ngwompo RF & Tilley DG. (2012). Development of a cardiopulmonary mathematical model incorporating a baro-chemoreceptor reflex control system. *Proceedings of the Institution of Mechanical Engineers Part H, Journal of engineering in medicine* **226,** 787-803.

Ursino M, Coli L, Brighenti C, Chiari L, de Pascalis A & Avanzolini G. (2000). Prediction of solute kinetics, acid-base status, and blood volume changes during profiled hemodialysis. *Ann Biomed Eng* **28,** 204-216.
